# Solid Tumors Pan Cancer Transcriptome: Tissue/Cancer specific expression groups at the Isoform-Level

**DOI:** 10.64898/2026.05.04.722705

**Authors:** Pallavi Surana, Matthew Obusan, Ramana V. Davuluri

## Abstract

Most of the human genome is transcribed into diverse isoforms whose tissue specificity is profoundly disrupted in cancer, yet isoform-level dysregulation remains poorly characterized across solid tumors. Here, we introduce STPCaT (Solid Tumors Pan-Cancer Transcriptome), an isoform-centric analysis extending TransTEx to systematically classify transcript expression across TCGA solid tumors and GTEx normal tissues. STPCaT reveals a striking collapse of normal tissue-specific programs in cancer, accompanied by the emergence of two dominant expression groups: cancer-high (Can_High_) and normal-high (Nor_High_) isoforms. We uncover a large repertoire of previously unannotated Cancer–Testis Antigens (CTAs), majority of which are absent from existing CTA databases, with broad relevance across multiple cancers, including gliomas. In pan-gliomas, consensus clustering and random-forest feature selection identify compact, highly discriminative isoform signatures that robustly stratify low-grade and glioblastomas with up to 97–98% accuracy using as few as five transcripts. These signatures recapitulate canonical glioma biology and highlight pathways linked to migration, development, and vesicle trafficking. Independent validation in the GLASS consortium cohort demonstrates cohort-specific trends that partially recapitulate primary findings, reflecting known biological heterogeneity across patient populations. Together, STPCaT provides a scalable, isoform-resolved resource for tumor stratification, CTAs discovery, and precision oncology applications across solid tumors.

**Teaser:** STPCaT uncovers an isoform-level collapse of tissue specificity across solid tumors, revealing a hidden landscape of diagnostic biomarkers and unannotated cancer-testis antigens.

## Introduction

The remarkable functional diversity of human cells arises from a genome where merely 1.5% encodes proteins, with the vast non-coding regions still largely under explored ^1^. Despite sharing identical DNA, cells achieve functional specialization through precisely regulated gene networks driving tissue-specific (TSp) expression patterns ^2,3^. Recent research reveals that primary transcription rather than alternative splicing as the dominant driver of cellular specialization, with thousands of genes exhibiting tissue-preferential expression profiles ^3-7^. The human genome, comprising approximately 50,000 genes: both coding and non-coding ^8-10^, generates an estimated 200,000 transcript variants that include roughly 100,000 protein isoforms ^11^. These isoforms exert substantial influence on gene expression through diverse post-transcriptional and epigenetic regulatory mechanisms, in TSp contexts ^12-15^.

Malignant transformation of cells and tissues fundamentally disrupts these carefully orchestrated expression patterns and their effectors. Cancer cells exhibit substantially more aberrant splicing events compared to paired normal tissues, with approximately 80% of these events not annotated in reference databases such as GENCODE ^16^. The functional consequences are profound; cancer cells frequently manipulate regulatory mechanisms to produce specific isoforms that facilitate drug resistance and enhance survival capacity ^11,17^. Notably, isoform switches leading to loss of DNA sequences encoding protein domains occur more frequently than expected, particularly in pan-cancer analyses, with several such switches identified as powerful biomarkers ^14,15^. About 58% of aberrant isoforms result from promoter switching events ^18^, highlighting the complex regulatory mechanisms underlying isoform diversity in cancer. These findings emphasize the need to investigate transcriptional profiles at the isoform level rather than solely at the gene level.

The clinical significance of isoform-specific analysis is well-established. Multiple studies have demonstrated that isoform-level expression profiles provide superior cancer signatures compared to gene-level expression analysis ^19,20^. Isoform-level gene signatures enhance prognostic stratification and improve cancer subtype classification accuracy ^21,22^. These observations align with the growing recognition of tissue specificity as a critical factor in disease initiation and progression ^23^. Despite this recognized importance, current therapeutic development strategies frequently neglect isoform diversity, with approximately 76% of drugs either overlooking potential target isoforms or targeting isoforms with variable expression across normal tissues ^11^.

Recent methodological advances have begun addressing the challenge of characterizing TSp expression at the isoform level. The development of resources such as TransTEx (Transcriptome Tissue Expression) and SRTdb have established frameworks for identifying TSp transcript level expression patterns ^5,13^. However, comprehensive isoform-level classification across multiple tumor types has remained challenging. While previous work demonstrated the utility of tumor-specific splice variants as cancer biomarkers across 10,000 specimens representing 33 cancer types ^9^, and implementing long read data to understand cancer transcriptome ^24^, a systematic framework for comparing isoform expression diversity across solid cancers with their relevance to the normal tissues is inadequate.

On performing differential expression analyses using edgeR v4 ^25^, comparing TCGA Solid Tumors to GTEx Normal tissues from UCSC Xena portal, suggest that tissue specificity is lost during cancer progression (Error! Reference source not found.). We observe that the tissue corresponding to the cancer counterpart show significant downregulation of TSp transcripts and non-TSp transcripts are upregulated in cancer. Null group attributes less than 3% of overall Null identified transcripts in GTEx. Further, those transcripts expressed in multiple tissues like Low and Wide are distributed across both groups. TEn group is more upregulated in UCS, UCEC, PAAD and others whereas, LUAD, LUSC, Gliomas are more downregulated, suggesting different patterns across cancers. These observations are consistent with previous studies showing that TSp gene expression patterns are often diminished or dysregulated during malignant transformation at the cancer site ^23,26-28^. This highlights the need to explore isoform-based expression in pan-solid tumors and their corresponding normal tissues. Identifying these expression patterns can help reveal biomarkers and different expression groups, which may further lead to the identification of actionable therapeutic targets.

To address these challenges, we developed <u>S</u>olid <u>T</u>umors <u>P</u>an-<u>Ca</u>ncer <u>T</u>ranscriptome (STPCaT) framework, which extends the TransTEx ^13^ methodology to categorize cancer (solid tumors) transcript expression profiles into Cancer (Can) -Specific (Sp), -Enhanced (En), -Widespread (Wide), Null and Low expression groups. We further evaluate their concordance to their normal matched tissues. Through systematic classification of transcripts into distinct expression groups across diverse Solid tumor types, STPCaT establishes a resource capable of identifying TSp and CanSp expression patterns. This framework enables clustering between different solid tumor types based on their isoform expression profiles and reveals molecular similarities among histologically or anatomically related cancer types, especially in Pan-gliomas.

Our approach identifies signature isoform patterns that not only distinguish between cancer types but also reveal shared molecular mechanisms with therapeutic potential. By focusing on isoforms with different expression patterns, we find groups like Cancer testis antigens (CTAs) and those that gain expression in cancer and lose in cancer compared to normal solid tissues. This work has relevance for precision oncology, as CanSp isoforms represent ideal diagnostic markers and therapeutic targets with potentially reduced off-target effects^5,9,29^.

In summary, this study presents a comprehensive characterization of transcript/isoform expression patterns across normal and cancer samples, establishing a framework for distinguishing between solid tumor types and identifying therapeutic candidates with potential clinical applications. By leveraging isoform-level resolution, we provide insights into cancer biology that may ultimately translate into improved diagnostic and therapeutic strategies for precision oncology.

## Results

### Global landscape of STPCaT: Comparison of Normal and Solid Tumor TransTEx

STPCaT is built using the different expression groups between Normal tissues from GTEx and the solid tumors from TCGA. We use mRNA isoform expression matrix and run the TransTEx algorithm separately on both the datasets. The algorithm considers 3 thresholds as summarized in **Fig. 1** (left panel). Using the isoform groupings from cancer and normal datasets we explore the global landscape and concordance between cancer and tissue specificity. We highlight 2 groups: Can_High_ which is highly expressed in cancer (1 or many solid tumors) and lowly expression in normals (Null/Low), and Nor_High_ which is the opposite (**Fig. 1** right panel). We explore the Can_High_ as potential oncogenic drivers and Nor_High_ as potential tumor suppressors. Aside from biomarker potential, we address the clinical relevance of CTAs and other expression groups in Pan-gliomas.

**Fig. 1:**
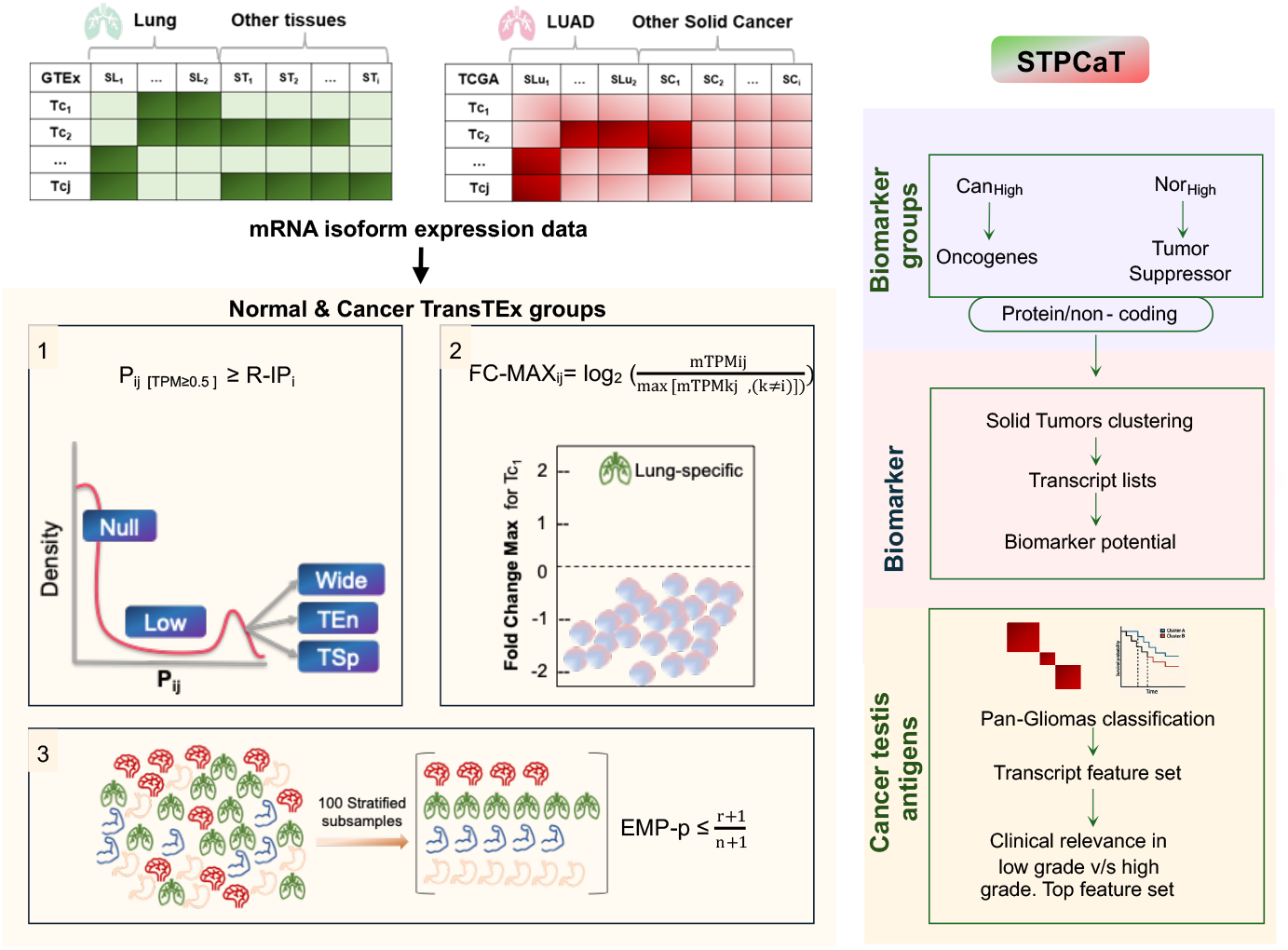
STPCaT resource. Utilize mRNA isoform expression data matrix from TCGA and GTEx consortiums (TransTExdb) are used to build STPCaT. The approach TransTEx takes to group the isoforms of any tissue or in this case of any cancer is shown in the left panel of the plot. Using these we propose STPCaT which identifies Pathways dysregulated in cancer gain or loss of function. Further, we identify biomarker groups, disrupted protein targets which could be used in therapeutics and the relevance of Cancer testis antigens in gliomas.

In STPCaT, we observe notable differences in transcript expression patterns. The diagonal in **Fig. 2A** shows minimal concordance between cancer and normal, with TSp and TEn each contributing less than 1%, while categories like Wide ∼6%, Null and Low >15% dominate. Additionally, TSp transcripts in normal tissues are largely reclassified as CanLow and CanNull, each comprising ∼4% of all transcripts. Two major patterns emerge from this comparison: Can_High_and Nor_High_ in cancer. Can_High_ events account for ∼5% and Nor_High_ for ∼23% of the common transcriptome, reflecting groups with significant divergence between cancer and normal. Further, more potential tumor suppressor transcripts lose expression in cancer. We highlight 2 more groups: Can_High_^Multiple^ for Can_High_ across multiple/all cancers and Nor_High_^Multiple^ for Nor_High_ across multiple/all cancers. Can_High_^Multiple^contributed to ∼1% of STPCaT whereas the other group to ∼12%. Cancers such as GBM, READ, and OV show high numbers of Can_High_transcripts, consistent with widespread transcriptional dysregulation in these tumor types (**Error! Reference source not found**.).

**Fig. 2:**
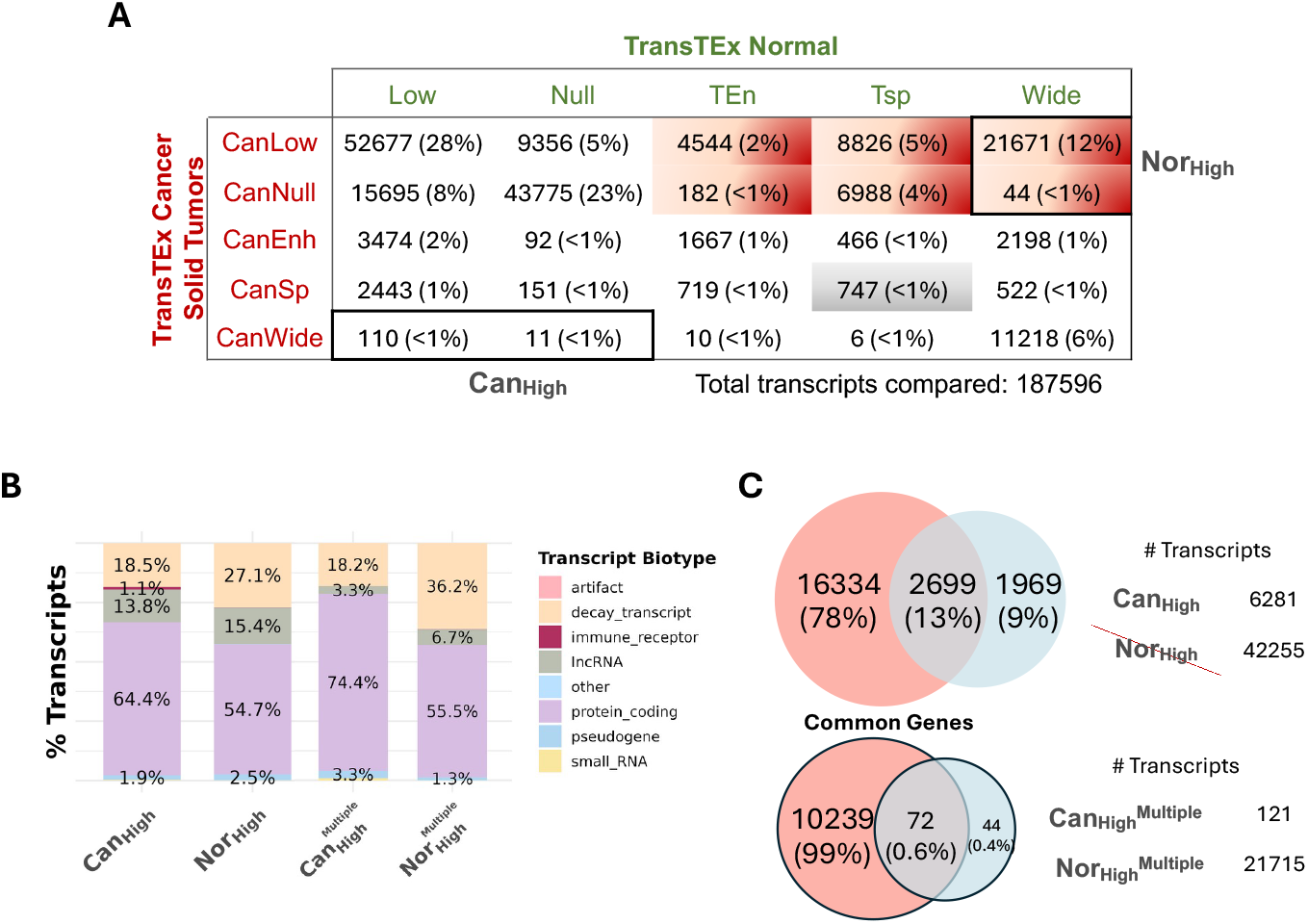
Global landscape of Cancer and Normal expression groups. (A) Concordance table between intersections and uncommon transcripts between expression groups in Cancer TransTEx and Normal adult tissue TransTEx where blue represents Can_High_(highly expressed in Cancer and minimally expressed in normal) and red represents Nor_High_(low/no expression in cancer but highly expressed in normal), grey box are for those TSp transcripts which become CanSp, and blue colored black outlined box is Can_High_^Multiple^ for Can_High_ across multiple/all cancers and red black outlined box is Nor_High_^Multiple^ for Nor_High_across multiple/all cancers. (B) Highlighting differences between transcript biotypes groups of Nor_High_, Can_High_, Can_High_^Multiple^ and Nor_High_^Multiple.^ (C) Common genes between the groups to identify the status of the alternate isoforms used between 2 distinct groups.

We next examined the transcript biotypes within the Can_High_ and Nor_High_ groups shown in **Fig. 2B**. Can_High_ transcripts were predominantly protein-coding, with a 10% higher representation compared to the Nor_High_ group. In contrast, Nor_High_ transcripts showed enrichment for non-coding biotypes, including a ∼2% increase in lncRNAs and ∼9% more decay-prone transcripts, indicating potential splicing-related disruptions. Both groups also contained a small proportion of pseudogenes. Within the ^Multiple^ categories we see ∼20% less protein coding transcripts in Nor_High_^Multiple^ and ∼18% increase in decay transcripts, indicating a stronger contribution from non-coding or unstable RNA isoforms. The Nor_High_^Multiple^ group comprises transcripts that are essential in normal tissues but lose expression in cancer, suggesting they must be rapidly degraded to prevent the production of truncated/deleterious proteins, yet the opposite pattern emerges in cancer. Less than 1% of genes overlapped between the ^Multiple^ groups, whereas 13% overlapped between Can_High_ and Nor_High_ (**Fig. 2C)**.

To study the role of different groups of expression related categories we test clustering 30 solid tumors in TCGA (Short forms of cancers in **Error! Reference source not found**.). We found that the CanSp, CanEn + TSp, TEn groups comprising ∼3.5k transcripts (∼75% of all transcripts in these categories), gave the best consensus hierarchical clustering results (k =30) when ranked by Mean Absolute Deviation (MAD). As an unbiased control, we repeated the analysis using the entire expression matrix and observed the optimal performance when selecting the top 15% of transcripts by MAD, corresponding to approximately ∼28k transcripts (**Error! Reference source not found**.). It is clearly observed that the lesser transcripts in the CanSpEn + TSpTEn group leads to more refined clustering of groups compared to overall transcripts which uses >10x the transcripts to cluster groups of the solid tumors. For instance, 23 LUSC samples mix with kidney cancer, HNSC cluster whereas they do not in the other group (**Error! Reference source not found.A-B)**.

### Landscape of testis specific transcripts in non-testis cancer

Among the testis-specific transcripts identified in the TransTEx, ∼14% (1,854 transcripts from 720 genes) were enriched as CTAs, referred to as the TestisSp_CTA group. Among the Cancer expression groups, out of the same 12,785 testis-specific transcripts (7,820 genes), only 15% (193 transcripts from 182 genes) were also classified as CanSp forming the Testis_CanSp_CTA group. Of these, 43 transcripts (38 genes) overlapped with known CTAs. In the Testis_CanSp_CTA group of transcripts we find majority (71) of the transcripts which are testicular cancer specific but some switch (105) into non-testis cancer majority of which are gliomas and OV (**Fig. 3A**).

**Fig. 3.**
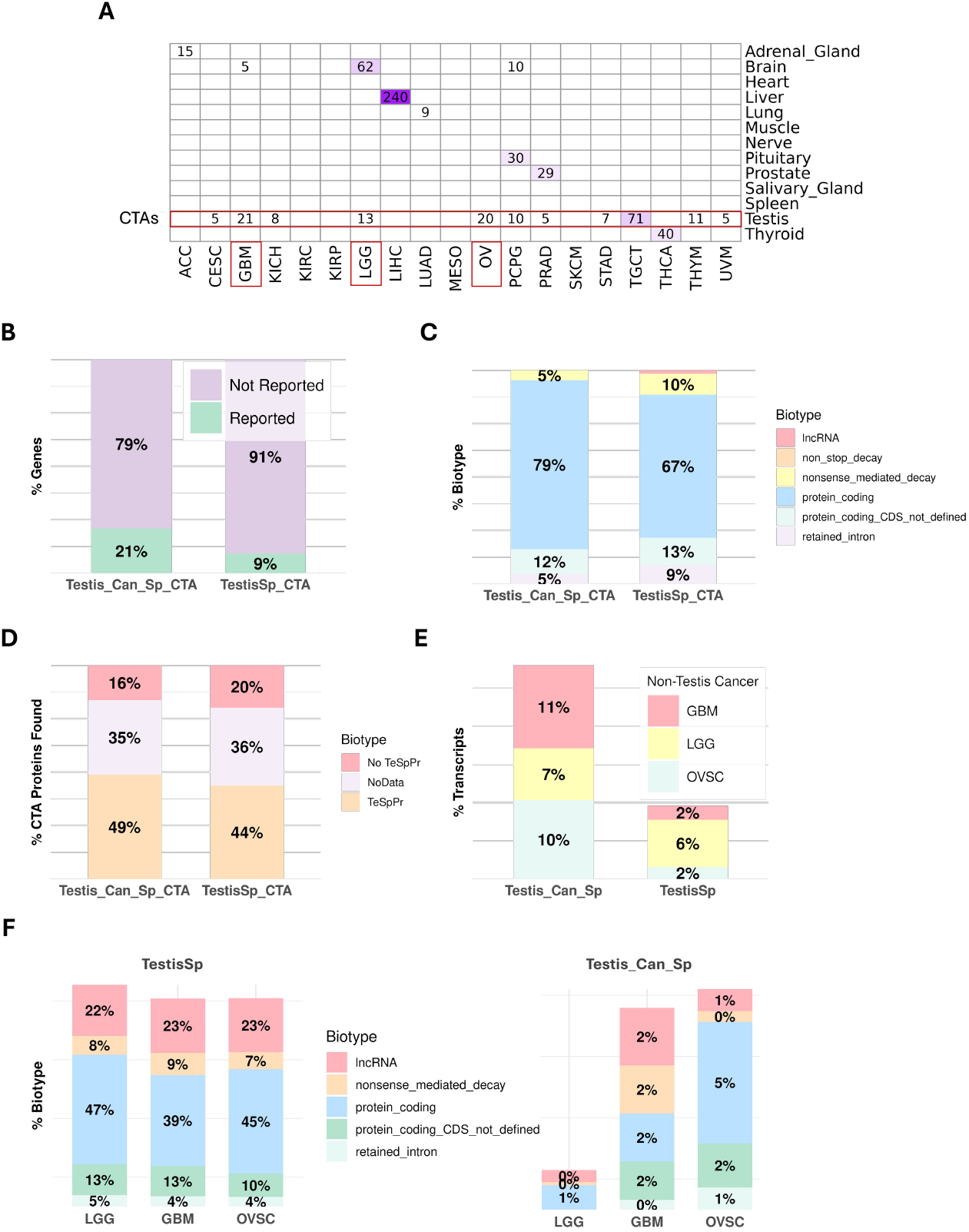
Cancer and tissue specific groupings of transcripts in STPCaT. (A) Top tissue and solid tumors with the number of transcripts in the boxes. Testis is highlighted in red box as many testis specific transcripts are expressed in cancers especially in Gliomas and Ovarian cancer in numbers >10. We explore this further in the direction of Cancer Testis Antigens (CTAs). **Note**: The subplots ahead are explored between 2 transcript groups: i) Testis-specific + CanSp (193 transcripts) and ii) All testis specific transcripts from TransTEx. Transcript groups i) and ii): B) mapped genes are queried against existing CTAs databases, and we highlight those reported v/s not. (C) transcript biotypes distribution. (D) proteins found in the CTAs databases highlighting the ones that are TeSpPr (testis specific protein) v/s not. (E) in non-testicular cancers like brain cancers (LGG, GBM) and ovarian cancer (OVSC). (F) transcript biotypes in brain cancers (LGG, GBM) and ovarian cancer (OVSC).

To study the biological relevance of STPCaT-identified CTAs, (**Fig. 3B** shows that a majority of TestisSp_CTA (91%) and Testis_CanSp_CTA (79%) transcripts have not been previously reported in existing CTA databases, highlighting their potential as novel immunotherapy targets. As shown in (**Fig. 3C**, most of these transcripts are protein-coding, with non-coding biotypes such as lncRNAs and decay-prone transcripts representing minor proportions (<5%). Protein classification (**Fig. 3D**) indicates that a large fraction of annotated entries belongs to the Testis-Specific Protein (TeSpPr) category, although many remain unannotated.

Next, we examine the distribution of these transcripts across three non-testis cancer types like gliomas (GBM, LGG) and OVSC (**Fig. 3E-F)**. Despite the relatively small number of Testis_CanSp transcripts (n = 193), they are detectable across all three cancers, with GBM showing the highest representation (**Fig. 3E**). Biotype distributions reaffirm that protein-coding transcripts dominate, though each cancer shows small (∼20%) but, consistent proportion of lncRNAs and retained introns. Collectively, these findings emphasize the isoform-level resolution of STPCaT in identifying previously unreported CTAs with relevance across multiple cancers, laying the foundation for future functional validation and therapeutic development.

Investigating the 34 CTAs that show up in gliomas (**Error! Reference source not found**.) we examined their biotypes and found lncRNAs enriched. Among these, 2 genes in GBM and 3 genes in LGG showed significant survival associations based on the TCGA Survival database (https://tcga-survival.com/). These genes include *ABCB9, RSPRY1, HMX1, KIF3A*, and *ERBB2*, each previously implicated in key aspects of glioma biology, such as therapy response, *SRP1*-dependent splicing promoting the mesenchymal phenotype, immune infiltration, and proliferative signaling ^30-34^.

### Cancer Testis antigens linked to Pan-Gliomas

We applied consensus hierarchical clustering to identify transcript sets that yield stable and biologically meaningful separation between lower-grade gliomas (LGG) and glioblastomas (GBM). Focusing on solutions dominated by two clusters (k = 2), we aimed to recover transcript signatures that maximize stability while minimizing misclassification. We use the WHO 2021 based classification of Across transcript categories, the most robust clustering performance was observed for transcripts that are expressed in cancer but not ubiquitously across cancers, followed by cancer-high (CanHigh) and cancer–testis antigens (CTAs) (Error! Reference source not found.).

Further analysis prioritized two transcript groups: CanSp_Enh_Wide_NorTSpTEnNullLow (495 transcripts), Can_High_ (628 transcripts) and TestisBrainSp_GBM_LGGsp (101 transcripts) (Error! Reference source not found. top, **Error! Reference source not found**.). These sets produced highly stable clustering, characterized by ≤10 transcripts with negative silhouette width and ≤2 misclassified GBM samples. Similarly, for k=3 we find that CanNull_NorTSpTEn group ith 717 transcripts best 3 groups classification into ACM, ODG and GBM of pan-gliomas (Error! Reference source not found. bottom). This group could potentially be significant tumor suppressor genes as they are highly specific and expressed in normals but completely lose expression in cancer. We show the distribution of the k=2 and k=3 clusters as well as the samples that are misclassified (**Fig. 4A, Fig. S7A)**. In the further analysis, we remove the misclassified samples from the clusters as well as the samples with negative silhouette width.

**Fig. 4.**
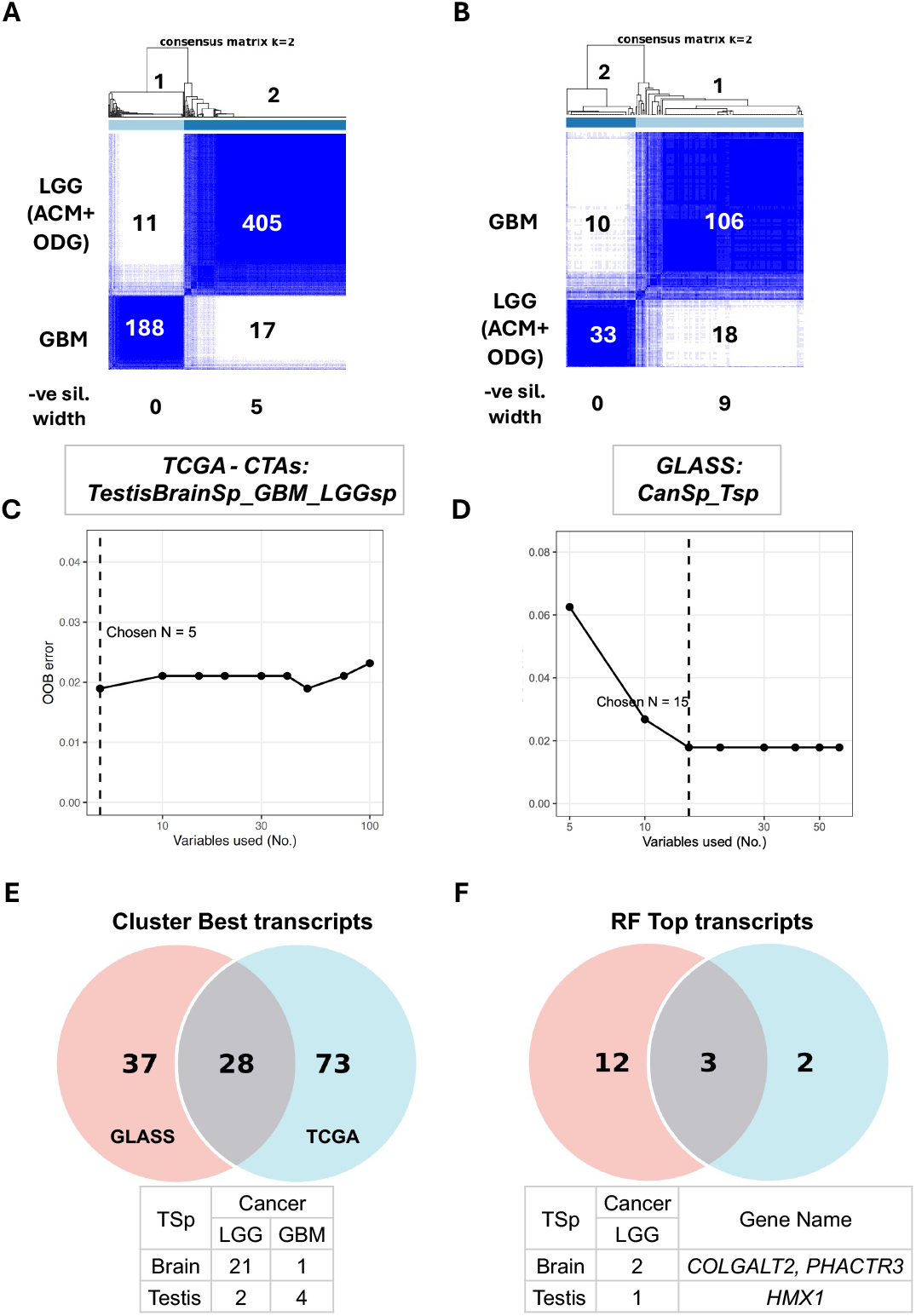
Consensus hierarchical clustering results for Pan-gliomas which includes GBM vs LGG where LGG includes ACM and ODG: (A) TCGA dataset with best result using testis and brain specific which are also specific to GBM and LGG, using all transcripts. (B) GLASS dataset without TCGA best result using specific transcripts in both cancer and normal tissues at 10% MAD. Random forest classifier results with top features, OOB error vs number of features of the train set plotted: (C) TCGA dataset with chosen N as low as 5. (D) GLASS dataset with chosen N as low as 15. Intersection of transcripts between the best results for k=2 between TCGA and GLASS datasets for gliomas: (E) Consensus hierarchical clustering best results comparison of the top transcripts in GLASS (65) and TCGA transcripts (101). (F) Random forest classifier top transcripts from GLASS (15) vs TCGA (5).

Random forest–based feature selection applied to these clusters yielded high classification performance (97–98% accuracy for both k=2 and k=3). Notably, a minimal subset of five transcripts comprising one testis-specific transcript and four brain-specific, LGG-specific transcripts was sufficient to achieve approximately 97% balanced test accuracy, highlighting the presence of a compact, highly discriminative transcript signature to classify GBM from LGG (k=2). We explore the genes that map to these transcripts (Error! Reference source not found.) and find convergence on key oncogenic processes, including cell migration and cytoskeletal regulation (*ARHGEF7, PHACTR3*), developmental reprogramming (*HMX1*), extracellular matrix remodeling (*COLGALT2*), and vesicle trafficking and secretion (*RAB6B*) **Fig. 4C**. Further, for k=3 we find top features as 50 transcripts (**Fig. S7B**).

When extending to k = 3, approximately 30 transcripts were consistently selected across iterations, of which 29 were tissue-specific, predominantly testis-specific transcripts, and one was a testis-enriched transcript. Among these, signaling components such as *LPAR* family members, central to lysophosphatidic acid–mediated glioma proliferation, migration, and invasion, and *MAPK10* (*JNK3*), a neural stress-activated kinase implicated in glioma cell survival and differentiation, emerged as recurrent features (Error! Reference source not found.). Together, these results indicate that highly restricted tissue-specific transcripts capture core biological differences between LGG and GBM, enabling robust clustering and accurate classification with remarkably small feature sets.

The distribution of canonical glioma molecular alterations across clusters was highly concordant with established disease biology and we explore this for both k=2 in **Error! Reference source not found.A** and k=3 in **Error! Reference source not found.B**. Cluster 1 (GBM-enriched) was characterized by canonical GBM alterations, including IDH wild-type status, frequent chromosome 7 gain with chromosome 10 loss, and TERT promoter mutations, with most tumors being ATRX-, DAXX-, and BRAF V600E–wild-type. MGMT promoter methylation was heterogeneous, consistent with known GBM variability. Cluster 2 (LGG-enriched) was enriched for IDH-mutant tumors, including both 1p/19q-codeleted and non-codeleted subtypes, and showed frequent ATRX alterations, MGMT promoter methylation, and absence of chromosome 7 gain/10 loss. BRAF alterations were rare and confined to LGG, while TERT promoter mutations were uncommon, reflecting lower-grade glioma biology. We observe similar patterns across the k=3 group in **Error! Reference source not found.B** recapitulates known taxonomy. Overall the classifiers are a useful tool with as low as 5 transcripts to classify GBMs vs LGG and ∼30 transcripts for ACM, ODG vs GBMs.

To validate the TCGA-derived glioma isoform clustering signatures in an independent cohort, we applied consensus clustering to the GLASS consortium dataset. For the two-cluster solution, the CanSp and NorSp transcript group (CanSp + NorSp; 65 transcripts) yielded the best performance, selected based on the highest ARI while maintaining acceptable average silhouette width and cluster purity (**Fig. 4B)**. The TCGA-optimal transcript group: Testis- and Brain-Specific isoforms with GBM and LGG specificity, was found to be a subset of this GLASS-identified group, indicating convergent feature selection across the two independent datasets (**Fig. 4E-F)**.

Critically, the three-cluster solution in GLASS recovered the same transcript group at the same MAD filtering threshold, despite the GLASS dataset containing a smaller transcripts set in the expression matrix (180,253 and 198,619 transcripts in TCGA) and an inverted histological composition (∼3x more GBMs than LGGs, compared to ∼2x more LGGs than GBMs in TCGA), supporting the replicability of these isoform signatures in an unseen cohort (**Fig. S7A-C, Table S7)**. The two-cluster separation in GLASS was less distinct than in TCGA, with a comparable number of misclassified samples (**Fig. 4A, B)**. In the three-cluster setting, GBMs formed two predominant subclusters while ODG and ACM co-clustered, with no independent ACM cluster resolved (**Fig. S7C**). Testing an alternative transcript group (CanEnGBM_LGG_TSpAll; 40% MAD) improved cluster purity but at the cost of average silhouette width (0.7664), and GBMs continued to bifurcate while ACM remained unresolved, likely due to limited sample size within this cohort (**Fig. S7D**).

## Discussion

Our study reveals a previously unrecognized level of divergence between normal and cancer transcriptomes by leveraging isoform-level tissue specificity. TransTEx analysis shows that classic tissue-specific/enriched categories: TSp and TEn, almost disappear in cancer, each contributing less than 1% of transcripts. Instead, two dominant opposing groups emerge: Can_High_, representing transcripts selectively activated in cancer, and Nor_High_, representing those selectively expressed in normal tissues but lost in tumors. Together, these groups encompass nearly a quarter of the shared transcriptome, illustrating a major axis of transcriptome remodeling between normal and malignant states. This reorganization highlights the importance of studying expression at the isoform level, where shifts in splicing and transcript stability become detectable.

A key outcome of this work is the discovery of a large pool of previously unreported Cancer-Testis Antigens (CTAs). Over 90% of TestisSp_CTA and nearly 80% of Testis_CanSp_CTA transcripts have not been annotated in existing CTA databases, despite displaying strong tissue restriction and cancer activation: both hallmarks of ideal immunotherapy targets. These findings demonstrate that STPCaT identifies a far broader and richer CTA landscape than gene-level methods, underscoring its potential to accelerate neoantigen discovery and CAR-T target development. The presence of these isoforms in multiple cancers, including gliomas, ovarian cancer, and testicular cancer, further supports their biological relevance.

Among gliomas, the identification of 34 CTAs with strong biotype diversity and survival associations reveals an unexpected link between testis-associated transcription and glioma biology. Notably, five genes namely, *ABCB9, RSPRY1, HMX1, KIF3A*, and *ERBB2* show significant survival differences in GBM or LGG, each tied to known pathways such as therapy resistance, mesenchymal transition, immune infiltration, and proliferative signaling. These results motivated us to explore the functional importance of CTA-expressing isoforms in gliomas. Using the WHO 2021 glioma classification, we show that a focused set of 101 STPCaT transcripts from the testis and brain specific + GBM and LGG-specific groups can robustly stratify gliomas into two stable groups with high silhouette width and cluster purity in the TCGA cohort. Strikingly, this small biologically structured panel outperforms genome-wide transcript selections. Further, as low as 5 transcripts are used by the classifier to distinguish these 2 major groups of pan-gliomas These transcripts exhibit minimal to no expression in cancer but are specific to and enhanced in normal adult human tissues, suggesting that the loss of normal tissue identity is a key feature of these tumors.

The clinical relevance of these clusters is further supported by misclassification analysis, which revealed minimal or no survival differences between samples misassigned as GBM versus LGG suggesting they could be indeed low-grade gliomas. This indicates that the STPCaT signature group captures meaningful biological variation rather than noise. Age, but not gender or race, was the only demographic factor associated with incorrect assignments, aligning with the known epidemiology of GBM and reinforcing the biological validity of our clusters. We further validated these robust groupings using related biomarkers in an independent cohort from the GLASS consortium. We observed comparable results for the 2-way groupings, despite the TCGA feature set being an incomplete subset of the GLASS dataset. Some observed dissimilarities are likely attributable to the inverted histological composition between the two datasets. However, the 3-way groupings could not be replicated in the GLASS cohort due to the limited sample sizes of the low-grade glioma subgroups within that dataset.

Collectively, these findings position STPCaT as a powerful analysis framework for identifying cancer-relevant isoforms, discovering novel CTA candidates, and enhancing tumor stratification with minimal yet biologically coherent transcript sets. Future work integrating proteomics, HLA-binding prediction, and functional perturbation studies will be critical to validating these isoforms as therapeutic targets. This analysis can be extended to single-cell RNA-seq data will be critical to determine if the “isoform expression collapse” occurs uniformly across the tumor bulk tissue or is driven by specific malignant subpopulations.

## Materials and Methods

### Datasets Used

We obtained transcript-level TPM expression data for TCGA ^35^ from the UCSC Xena platform ^36^, creating a comprehensive database termed Cancer TransTEx (CanTransTEx). For consensus hierarchical clustering, we specifically used the transcript-level log2(expected_count + 1) data, encompassing 198,619 transcripts across 9,805 samples. Our analyses were restricted to solid tumor samples only, comprising 8,977 samples spanning 30 cancer types. For Normal TransTEx, we adopted the tissue-specific groupings provided by TransTExdb: https://bmi.cewit.stonybrook.edu/transtexdb/. Clinical annotations for TCGA samples were also retrieved from UCSC Xena.

### Application of TransTEx

We utilized the TransTEx R package (v 0.1.0) to classify transcripts or genes into five distinct expression groups based on their expression probabilities across tissues. This process involves calculating three key metrics for each transcript: FC-MAXij (maximum fold change), EMP-p (subsampling-based empirical p-value), and the probability estimate Pij. Thresholds for classification were determined using right (R-IPi) and left (L-IPi) inflection points from expression distributions.

Transcripts were grouped as follows:

1. Determine the number of tissues (k) for which the probability of transcript expression is above the R-IPi threshold (Pij ≥ R-IPi).
2. Classify transcripts based on k:
  - Low or No Expression (Low transcripts): k = 0.
  - Tissue-Specific Expression (TSp transcripts): k = 1.
  - Tissue-Enhanced Expression (TEn transcripts): k = 2 to 50% of tissues.
  - Widespread Expression (Wide transcripts): k > 50% of tissues (14-total tissues).
3. For TSp and TEn transcripts, FC-MAXij and EMP-p values were computed, and transcripts with FC-MAXij ≥ 1 and EMP-p ≤ 0.05 were retained in their respective classes.
4. For transcripts where k = 0, classification as either Null Expression or Low Expression was determined by comparing their probabilities to the L-IPi cutoff (left peak cutoff). Transcripts with probabilities below L-IPi in all tissues were grouped as Null Expression; otherwise, they were classified as Low Expression ^13^

We have summarized the pipeline of the same in **Figure 1** where we used the transcript mapping generated by TransTExdb as well as implement CanTransTEx.

### Comparison with Cancer Testis Antigens

We consolidated our Cancer Testis Antigen (CTA) reference set by integrating gene lists from multiple curated sources and studies: the Cancer Testis Antigen Burden (CTAB) metric outlined by ^37^, the tumor CTA catalogue by ^38^, the curated CTDatabase repository ^39^, and the genomic-instability-associated CTA list from ^40^. Gene identifiers were extracted directly from supplementary data or online repositories provided by these studies. Subsequently, these lists were merged and deduplicated to yield a unified, non-redundant CTA reference gene panel. We compared transcript-gene pairs from our analyses against this comprehensive CTA panel to identify enriched and relevant CTA-associated transcripts.

### Consensus clustering analysis

Consensus clustering was performed following methods recommended by ^19,21^. We downloaded the isoform-level, log2-normalized expected counts matrix for TCGA from UCSC Xena ^36,41^. For The Glioma Longitudinal Analysis Consortium (GLASS) we downloaded the data from the current release hosted on Synapse (syn26465623) ^42^. To ensure comparability with primary tumor biology and to avoid overlap with TCGA samples, only primary tumors denoted by the “TP” in sample IDs were used, and any TCGA-derived samples were excluded. This cohort was not used in the comparative STPCaT analysis and therefore represents a fully unseen validation dataset. For both datasets, lowly expressed transcripts were filtered out, and the remaining transcripts were ranked based on mean absolute deviation (MAD) across the sample cohort, retaining the most variable features. The resulting filtered matrix was normalized using median-MAD normalization with the data.normalization function (type=“feature_Median”) in the CancerSubtypes R package (v1.17.1). Hierarchical consensus clustering was then conducted using Pearson distance metrics and 50 resampling iterations for cluster numbers ranging from K = 2 to K = 10 (for Subtyping) or K=30 (for Pan-Cancer). Cluster quality and optimal cluster selection were determined using cumulative distribution function (CDF) plots, consensus delta-area plots, silhouette width analysis (via factoextra package v1.0.7), and, where reference labels were available, the Adjusted Rand Index (ARI).

### Survival and Pathway Analysis

Survival analysis was conducted using the survival package (v3.8.3) in R ^43^. A univariate Cox proportional hazards regression model was applied to assess the relationship between isoform-level transcript expression and patient survival outcomes. Hazard ratios (HR), confidence intervals, and statistical significance (p-values) were computed, and results were corrected for multiple hypothesis testing using the Benjamini–Hochberg method to calculate false discovery rates (FDR) ^44^. Significant survival-associated transcripts were identified based on an FDR threshold of 0.05 and further classified as risk (HR >= 1.5) or protective (HR <= 1/1.5). Pathway enrichment analysis was performed using the tool cGSA. We specifically utilized the gene set libraries for pathway enrichment like KEGG, WikiPathways, MSigdb and Reactome.

### Top Transcript feature selection using Random Forests

Transcript-level features predictive of pan-glioma cluster membership were identified using a Random Forest–based stable feature selection approach. Transcripts were first filtered to the transcripts which gave the best consensus clustering results. Only samples with positive silhouette widths were included to ensure robust cluster assignments. Random Forest classifiers (1,000 trees) were trained over 50 iterations using random 80% subsamples of the data. Feature importance was assessed using permutation-based mean decrease in accuracy and Gini index. Transcript stability was determined by the frequency of appearance among the top 50 features across iterations, with final ranking based on stability frequency and mean importance. Selected transcripts were annotated for downstream biological interpretation.

## Acknowledgments

We would like to thank the Davuluri lab and computing servers at Stony Brook University to make this work possible.

## Funding

This work was supported by:

National Library of Medicine/National Institutes of Health [R01LM013722 to R.D.]

## Author contributions

Conceptualization: PS, RD

Methodology: PS, RD

Investigation: PS

Visualization: PS, MO

Supervision: RD

Writing—original draft: PS

Writing—review & editing: PS, MO, RD

## Data and materials availability

All data are available in the main text or the supplementary materials. We will make the entire STPCaT groupings available upon acceptance of the paper.

